# We are the sensors of consciousness! A review and analysis on how awakenings during sleep influence dream recall

**DOI:** 10.1101/2024.11.13.623388

**Authors:** Benjamin Stucky

## Abstract

**Purpose:** Since the 1930s, researchers have awakened people from different stages of sleep to record what they have experienced. While some aspects, including asking whether participants had dreams or thoughts before awakening, largely remain the same, others, such as the method of awakening, vary greatly. In addition, it is often assumed that the influence of participant characteristics, such as personality traits, motivation, memory, and attention, is reduced by collecting experiences immediately after they occur, rather than through delayed morning recall. However, the extent to which these variables influence dream recall upon awakening has not yet been thoroughly investigated.

**Materials and methods:** To explore possible contextual and individual influences, this review analyzed 69 awakening studies conducted between 2000 and 2024 and utilized the DREAM database. Differences between sleep stages were quantified and experiences analyzed using the categories ‘with recall’, ‘without recall’, and ‘no report’.

**Results:** Similar levels of null reports were found in NREM stage 2 and stage 3. Significant factors affecting dream recall included the method of awakening (lower recall with an alarm compared to calling the participant’s name), the number of study days (reduced recall for multiple days) and the sleep environment (higher recall at home compared to the laboratory), along with participant characteristics beyond age, sex and study design. Recall rates from NREM sleep are particularly sensitive to the method of awakening and interindividual differences.

**Conclusion:** Both the awakening procedure and participant characteristics influence the amount of reported sleep experiences, which can impact study outcomes, such as the identification of neural correlates of consciousness. Therefore, greater emphasis needs to be placed on how experiences are collected and on participant characteristics, such as openness to experience or familiarity with different states of consciousness.

**Plain Language Summary:** Have you ever wondered how often you dream while you sleep? We reviewed 69 studies where people were woken up to share their experiences. They reported dreaming most of the time, even in deep sleep. You might ask, “Why don’t I remember much in the morning then?” Well, maybe you forgot your dream because a lot of time has passed. These studies capture the experience right after it happens, but even then, some people remember dreams more than others. This could be because they are more focused on their dreams. How you wake up also matters. If someone shakes you awake, you may forget your dream, but if they gently call your name, you have a better chance of remembering it. You might also fail to recognize more subtle experiences, such as a sense of calm. So perhaps sleep is not as unconscious as we thought, leaving much to be explored in future studies.

## Introduction

### Early beginnings

In 1937, not long after the first human electroencephalogram (EEG) recording,^1^ Loomis, Harvey and Hobart published their pioneering work on classifying sleep stages.^2^ In that same paper, they collected an experience report after a doorknob twist woke up a study participant. They assumed that the sound was incorporated into the participants’ dream, since the experiential report read: “someone had made a noise like a train.” The doorknob twist happened when the participant was in what we now call non-REM (NREM) stage 3 (N3). Even though the authors attributed the dream to one minute of lighter sleep occurring between the sound and the collection of the experience report, it could mark the first confirmed NREM sleep experience.

Just one year later, another experiment by the same authors and Davis and Davis investigated multiple spontaneous and sometimes provoked sleep onset experiences.^3^ There we find a description of one of the earliest ‘white dreams’ from a provoked NREM stage 2 (N2) awakening. A ‘white dream’, or an experience without recall, is the impression of having just experienced something but not being able to remember any details. This work along with another landmark paper by Blake, Gerard, and Kleitman laid the groundwork for studies provoking multiple awakenings within a sleep episode.^4^

### REM sleep is dream sleep?

It wasn’t until the early 1950s that these types of laboratory based awakenings with subjective reports were taken up again in a rigorous manner.^5^ The seminal paper on REM sleep, characterized by wake-like EEG patterns and rapid eye movements with closed eyelids, suggested that dreams mainly occur in that stage.^6,7^ This finding was further consolidated by a large study with multiple participants, multiple nights and multiple awakenings per night that reported dreams in 80% of the REM sleep awakenings but only 7% from NREM sleep.^8^ Another paper by Dement and Wolpert^9^ used brief awakenings to associate eye movement directions with the reported gaze in the dream. All of this reinforced the view, still common in popular culture today, that dreams occur only in REM sleep.

But this idea was met with contrary evidence early on. In a study from 1959, participants who recalled dreams on most nights, so called high recallers, had on average 53% dream recall from NREM sleep compared to low recallers with 17%.^10^ Over time, accumulating evidence showed that recalling experiences from NREM sleep is not uncommon, finding experiences in up to 70% of the awakenings.^11^ Despite that, the idea of REM sleep being dream sleep did not suddenly disappear; instead, there were attempts to explain NREM dreams as occurring during preceding REM sleep or the awakening process itself.^12^ Such arguments were weakened by Foulkes who showed that dreams can be recalled in the absence of REM episodes.^11^ He aimed to capture a broader range of experiences by introducing the now ubiquitous question ‘What was going through your mind?’ instead of ‘What were you dreaming?’. Furthermore, he argued that differences in questions and the exclusion of less visual experiences might have contributed to inconsistent recall rates. In doing so, he redirected attention towards examining differences between NREM and REM experiences.^12,13^ Subsequently, it has been observed that REM reports tend to be longer and more vivid than NREM ones^14–19^, though without precise delineation.^16^

### Neural Correlates of Dreaming

As it became evident that all sleep stages contain experiences, the focus shifted towards identifying transient neurophysiological changes, either common or unique to NREM and REM sleep, that account for obtaining dream reports.^20,21^ In this vein, Crick and Koch rejuvenated the search for neural correlates of consciousness.^22–24^ They focused on ‘phenomenal consciousness’, the felt qualities of the present moment.^25^ This definition of consciousness differs from others, as some felt states might arise without cognitive processing,^26–28^ can go unnoticed, be forgotten, or lack bodily reactions.^29–31^ In this review we use ‘consciousness’, ‘dreaming’, and ‘experience’ interchangeably to mean phenomenal consciousness. The subjective reality challenges physicalism, raising the question why presumably ‘non-conscious’ physical processes lead to any felt sense, known as the hard problem.^32^ While not solving this mystery,^33^ Crick and Koch argued that correlating phenomenal consciousness with neural activity is a first step.^22^ Awakening studies providing subjective reports are well-suited for this purpose^34^ and have in part led to proposed laws of consciousness.^35–37^ High-density EEG studies point to the ‘posterior cortical hot zone’ as a correlate.^20,34,38,39^ However, according to a recent review, there is no consensus on the neural correlates of dreaming.^40^ A ‘dream catcher’ experiment involving separate data gathering and analysis teams, also failed to predict dreams from the EEG better than chance at various levels of blinding.^41^ This uncertainty is further emphasized by two studies attempting to correlate the diversity of dreams with EEG signal complexity, resulting in null or conflicting findings.^42,43^ This raises the question: where do these inconsistencies come from?

### Today’s missing ingredients

Ever since those initial investigations, some main ideas behind awakening protocols have largely remained consistent. This involves brief awakenings followed immediately by experience reports. Other aspects have varied considerably and their impact on dream recall remains largely unexplored.

For example some studies provoked multiple awakenings in early sleep, termed ‘Early-Night Serial Awakenings’.^44^ Another procedure, the ‘Serial Awakening Paradigm’, is characterized by frequent random awakenings during the whole night.^45^ The ‘Sleep Interruption Technique’ induces sleep onset REM or NREM through awakenings followed by brief periods of sleep and experience sampling.^46,47^ Not just night sleep has been under investigation, a multiple nap protocol with spontaneous awakenings as well as single nap protocols have been developed.^48–50^ Multiple awakenings have not been confined to the laboratory setting alone. Home-based awakenings utilized portable devices with algorithms to predict sleep stages^51–55^ and portable polysomnography (PSG) at home.^56,57^ There were also single night and multiple night studies.^58,59^ Questions about participants’ mental activity and dreams are the most common, with a minority asking slightly different questions,^46,51,60,61^ while many include secondary study-specific questions.^43,45,51,58,62^ Further variability can be found in the methods used to wake up participants, some call the participant by name^58,63–65^, use an alarm^45,56,66^, or sounds^41,47,62,67^, others directly ask the question, knock or enter the room.^52,68–71^ Other lines of advance include interventions prior to awakenings^72–77^, non-healthy subpopulations^56,57,62,71,78,79^ or pharmacological interventions.^16^

Adding to this diversity, there might be interindividual effects. It has been known for a while that experiences recalled during the night just after they happened can be forgotten by the same participant in the morning, highlighting the importance of memory.^80^ A recent paper found that attentional processes might even play a larger role in dream recall.^81^ Other participant characteristics associated with spontaneous morning dream recall include age, sex, attitude towards dreams, openness to experience, and creativity.^82–86^ It has been assumed that such effects are less pronounced for studies collecting the experience near to when it happens compared to delayed morning recall.^34,87^ But this hypothesis remains to be tested.

Given that many objectives in dream research, such as identifying neural correlates of dreaming, depend on subjective reports—which can be influenced by awakening procedures and participant characteristics—careful investigation into the effects of context and personal characteristics with a focus on awakening studies is warranted.

### In short

Awakening studies have a rich and interesting history and have been used to identify neural correlates of dreaming. But what is currently missing is an analysis on how contextual variables such as the sleep environment, the questions being asked, the awakening procedure, and participant characteristics influence dream recall. Consequences could be that research objectives depending on subjective reports and interpretations thereof get biased. To shed light on these effects, we review and analyze papers between the years 2000 – 2024 with awakenings from different sleep stages and subsequent experience sampling. For testing the presence of interindividual effects, we use the freely accessible DREAM database (v 4)^88^, consisting of participant level data from some of the recent laboratory based awakening studies.

## Material and methods

### Study inclusion criteria

We have conducted a non-systematic review of studies with awakenings from different stages of sleep to collect experiential reports. The resulting list of studies aims to capture many relevant studies, though it is not meant to be exhaustive. Inclusion criteria were:

1. Experiential reports are associated with at least one sleep stage, either N1, N2, N3, NREM or REM.
2. Percentages of experiences (recall rates) are available, or inferable, for at least one stage and one experience type (‘experience with recall’, ‘experience without recall’ or ‘no report’).
3. Participants are healthy.
4. No interventions are present.
5. The studies were published, or made accessible on an online repository between January 2000 and July 2024.
6. Recall rates from extensive subcategories, like many collection timepoints, were omitted to avoid disproportionately increasing the weight of individual studies.
7. Whenever studies shared the same data and reported the same recall rates, only one study was retained. Sometimes the data basis was only partially shared or the same data used but complementary information reported, in which case all were included.

The studies were mainly searched on Google Scholar (https://scholar.google.com/) and PubMed (https://pubmed.ncbi.nlm.nih.gov/) with the following keywords: *dream recall, dream reports, serial awakening(s), serial awakening paradigm, serial awakening sleep, sleep interruption technique, early-night serial awakenings, sleep interruption technique, laboratory dream, laboratory awakening dream, laboratory awakening sleep, awakening sleep, awakening dream, consciousness sleep*.

Additionally, some studies were found through the citation list of other studies and a small amount through queries into Microsoft Copilot, an AI assistant.

### Extracted Variables from Studies

Whenever available, we searched and extracted the following information within the main paper or the supplementary information of each study:

1. Percentages of experience with recall, experience without recall or no report. In cases where experience with and without recall were merged, the no report category was inferred (ie 100% - percentage of experience with and without recall).
2. The awakening method. For balanced analyses a factor with three levels was constructed: awakening with an alarm, buzzer, sound or tone, by calling the participant by name and a category including ‘other’ methods.
3. The question used for experience type categorization. The three main levels were ‘what went through your mind’, ‘what did you dream’ and ‘other’.
4. Number of awakenings per recording with levels ‘repeated’ or ‘single’.
5. The sleep environment with levels ‘laboratory’ or ‘at home’.
6. The type of sleep, if it was a nap or a night sleep.
7. The study days, consisting of multiple or single experimental days.
8. Participant characteristics like mean age and percentage of females, calculated per study or study subset. If only an age range was given, the mean age was approximated by the midpoint (ie maximum age – minimum age divided by two).
9. Number of unique participants per study or study subset.
10. Number of awakenings per study or study subset.
11. If the awakening was explicitly mentioned as being gentle.

Some values were not explicitly given, but could be inferred or approximated. Inferred values are indicated it in the supplement Table S1. Some factor levels consisted of a mixture of other levels or were uncommon, these received the label ‘both’ or ‘other’.

### Data Analysis: Review

All analyses were carried out with the programming language R (v 4.3.2)^89^ in the integrated development environment RStudio (v 2024.4.2.764).^90^

Three independent linear regression models were constructed, one for ‘experience with recall’, another for ‘experience without recall’ and lastly one for ‘no report’. In order to retain statistical power, we condensed all NREM sleep stages into a single level (consisting of NREM or N1, N2 and N3). Since only experience with recall had a substantial amount of data, we used it to construct a more extensive model with variables: stage, awakening method, question type, mean age, percent female, repeated awakenings per recording, sleep environment, study duration and sleep type. For these analyses the ‘other’ and ‘both’ levels were omitted. The models for experience without recall and no report only included sleep stage as an independent variable. The p-values were corrected for multiple comparisons with the Benjamini-Hochberg method per model.^91^ Additionally, comparisons between N2 and N3 were carried out with t-tests and were also corrected with Benjamini-Hochberg. All figures report the corrected p-values, whereas tables report both, the raw and corrected p-values. Cohen’s d values and corresponding effect size categories were additionally computed from the models by the emmeans (v 1.10.2)^92^ and effectsize (v 0.8.8)^93^ packages. The effect size categories were ‘negligible’ for absolute Cohen’s d values < 0.2, ‘small’ if 0.2 ≤ | d | < 0.5, ‘medium’ 0.5 ≤ | d | < 0.8, and ‘large’ if | d | ≥ 0.8.

The experience with recall model only included main effects. To check whether some interactions are important, we constructed a model with all interaction terms per sleep stage. A stepwise AIC model selection with both directions was carried out with the MASS package (v 7.3-60).^94^ The main effects were retained in each step. These analyses can be found in the supplement Table S2. Furthermore, the supplement contains analyses for potential effects of study year (Figure S1), number of awakenings per participant in single night studies (Table S3) and gentle awakenings (Figure S2) on percentages of different experience types.

### Data Analysis: DREAM database

For the DREAM database we only included healthy participants and sleep stages that were not labeled ‘Unknown’ or ‘#N/A’. Since the ‘Subject.ID’ variable was not fully unique between studies, we created a new unique identifier per participant and study. An additional age factor was constructed with levels less or equal than 30 years or more than 30 years of age. This cutoff point was chosen, since it corresponds to the rounded mean age of 29.5 amongst the studies.

The subset of studies containing either experience with recall, experience without recall or no report, were identified. For each such report type and sleep stage, we calculated participant-based percentages. To ensure meaningful estimates, we only kept those percentages that were based on 3 or more data points per person.

First, generalized mixed effect models with sleep stages as fixed effects and study identifier as random intercept were estimated. Since the distributions were bounded by and highly skewed towards 0% and 100%, generalized ordered beta mixed model^95^ with logit link function implemented in the glmmTMB package (v 1.1.9)^96^ were used. This model is specifically designed for bounded outcomes. The results were corrected for multiple testing with Benjamini-Hochberg and amended with Cohen’s d values and the same effect size categories as mentioned above. Additionally, to test for participant characteristics, we made use of permutation tests. Permutation tests have the advantage of making minimal assumptions on the data.^97^ The permutation test was constructed with the standard deviation of participant-based percentages as a test statistic. This was chosen, since it is a measure of how variable participants are between themselves. Thus, one can test if the observed variability is expected due to pure chance or rather due to participant grouping. Resampling of experience labels was done 10’000 times. Importantly, the structure of sex, age category, and study id was kept intact, meaning labels were randomly resampled only within the groupings but not between. This way one can control for these factors and the resulting significant effects are indicative of participant characteristics beyond sex, age and other variables of a given study. After each resampling the percentages per person were calculated again for NREM (with additionally keeping stage grouping—N1, N2, and N3—intact in permutations) and REM sleep. All permutation tests reported in the main paper have been corrected for multiple testing with Benjamini-Hochberg. To see if observed NREM effects hold true for a single sub-stage, we calculated the permutation test only on N2, which contained the largest amount of data. These additional analyses are reported in the supplement Figure S3.

## Results

### Descriptive statistics: Review

In total, 69 studies were found who met the inclusion criteria.^16,18,19,34,42–50,52,54,55,57–62,64–70,72,75,98–100,100–135^ All studies reported experience with recall percentages and 22 studies additionally captured experiences without recall and no report, see Table 1.

**Table 1.** Included studies and their characteristics. Here the 69 studies that were included in the review are listed and sorted by year of publication. In particular, the columns awakening method (method), question type (question), awakenings per recording (repeated), sleep environment (setting), type of sleep (sleep), and the study days (days) as well as the presence (yes) or absence (no) of reported percentages for experience with recall (w/ recall), experience without recall (w/o recall) or no report are presented.

| Publication | Year | Method | Question | Repeated | Setting | Sleep | Days | w/<br>recall | w/o<br>recall | No<br>report |
| --- | --- | --- | --- | --- | --- | --- | --- | --- | --- | --- |
| Parker et al.<sup>69</sup> | 2000 | other | other |  |  |  |  | yes | no | no |
| Takeuchi et al.<sup>47</sup> | 2001 | alarm | mind | repeated | lab | night | >1 | yes | yes | yes |
| Stickgold et al.<sup>55</sup> | 2001 | other | mind | repeated | home | night | >1 | yes | no | no |
| Takeuchi et al.<sup>46</sup> | 2003 | alarm | other | repeated | lab | night | >1 | yes | yes | yes |
| Palagnini et al.<sup>105</sup> | 2004 | name | mind | single | lab | nap | >1 | yes | no | no |
| Fosse et al.<sup>18</sup> | 2004 | other | mind | repeated | home | night | >1 | yes | no | no |
| Wittmann et al.<sup>65</sup> | 2004 | name | mind | repeated | lab | night | >1 | yes | yes | yes |
| St-Onge et al.<sup>100</sup> | 2005 | other | mind | repeated | lab | night | 1 | yes | no | no |
| Strauch<sup>112</sup> | 2005 | name | dream | repeated | lab | night | >1 | yes | no | no |
| Grenier et al.<sup>135</sup> | 2005 | name | mind | repeated | lab | night | 1 | yes | no | no |
| Wamsley et al.<sup>106</sup> | 2007 | name | mind | repeated | lab | night | 1 | yes | yes | yes |
| Weigand et al.<sup>67</sup> | 2007 | alarm | mind | both | lab | night | 1 | yes | no | no |
| Daoust et al.<sup>103</sup> | 2008 | other | mind | repeated | lab | night | 1 | yes | yes | yes |
| Wamsley et al.<sup>106</sup> | 2008 | name | mind | repeated | lab | nap | >1 | yes | no | no |
| Noreika et al.<sup>44</sup> | 2009 | alarm | mind | repeated | lab | night | >1 | yes | yes | yes |
| Lusignan et al.<sup>111</sup> | 2009 | other | mind | repeated | lab | night | >1 | yes | yes | yes |
| Lara-Carrasco et al.<sup>118</sup> | 2009 | alarm | mind | repeated | lab | night | 1 | yes | no | no |
| Lusignan et al.<sup>70</sup> | 2010 | other | mind | repeated | lab | night | 1 | yes | yes | yes |
| Wamsley et al.<sup>110</sup> | 2010 | other | mind | repeated | lab | nap | 1 | yes | no | no |
| Wamsley et al.<sup>54</sup> | 2010 | other | mind | repeated | home | night | >1 | yes | no | no |
| Schredl et al.<sup>126</sup> | 2010 | name | mind | repeated | lab | night | 1 | yes | no | no |
| Chellappa et al.<sup>50</sup> | 2011 | other | dream | single | lab | nap | >1 | yes | no | no |
| Kahan et al.<sup>51</sup> | 2011 | alarm | other | repeated | home | night | 1 | yes | no | no |
| Blagrove et al.<sup>130</sup> | 2011 | alarm | mind | repeated | lab | night | >1 | yes | no | no |
| Marzano et al.<sup>98</sup> | 2011 | name | dream | repeated | lab | night | 1 | yes | no | no |
| Oudiette et al.<sup>16</sup> | 2012 | name | mind | repeated | lab | night | >1 | yes | no | no |
| Kusse et al.<sup>119</sup> | 2012 |  | mind | repeated | lab | nap | >1 | yes | no | no |
| Chellappa et al.<sup>120</sup> | 2012 | other | dream | single | lab | nap | >1 | yes | no | no |
| Stenstrom et al.<sup>60</sup> | 2012 | name | other | repeated | lab | night | >1 | yes | yes | yes |
| Siclari et al.<sup>45</sup> | 2013 | alarm | mind | repeated | lab | night | >1 | yes | yes | yes |
| Schredl et al.<sup>104</sup> | 2014 | other | mind | repeated | lab | night | 1 | yes | no | no |
| Sikka et al.<sup>101</sup> | 2014 | alarm | dream | repeated | lab | night | >1 | yes | no | no |
| Yu<sup>99</sup> | 2014 |  | mind | repeated | lab | night | 1 | yes | no | no |
| Eichenlaub et al.<sup>68</sup> | 2014 | other | mind | single | lab | night | 1 | yes | yes | yes |
| Carr et al.<sup>108</sup> | 2015 | alarm | dream | single | lab | nap | 1 | yes | no | no |
| van Rijn et al.<sup>52</sup> | 2015 | alarm | mind | repeated | both | night | >1 | yes | no | no |
| Yu<sup>114</sup> | 2016 |  | mind | repeated | lab | night | 1 | yes | yes | yes |
| Nefjodov et al.<sup>127</sup> | 2016 | other | mind | repeated | lab | night | 1 | yes | no | no |
| Scarpelli et al.<sup>64</sup> | 2017 | name | mind | repeated | lab | nap | 1 | yes | yes | yes |
| Siclari et al.<sup>34</sup> | 2017 | alarm | mind | repeated | lab | night | >1 | yes | yes | yes |
| Sikka et al.<sup>121</sup> | 2017 | alarm | dream | repeated | lab | night | >1 | yes | no | no |
| Solomonova et al.<sup>48</sup> | 2018 | alarm |  | repeated | lab | nap | 1 | yes | no | no |
| Eichenlaub et al.<sup>116</sup> | 2018 | alarm | mind | repeated | lab | night | 1 | yes | no | no |
| Feige et al.<sup>62</sup> | 2018 | alarm | mind | repeated | lab | night | >1 | yes | no | no |
| Siclari et al.<sup>59</sup> | 2018 | alarm | mind | repeated | lab | night | >1 | yes | yes | yes |
| Blagrove et al.<sup>113</sup> | 2019 | alarm | mind | repeated | lab | night | 1 | yes | no | no |
| Wamsley et al.<sup>122</sup> | 2019 | name | mind | repeated | lab | night | 1 | yes | no | no |
| Schoch et al.<sup>123</sup> | 2019 | other | mind | repeated | lab | night | 1 | yes | no | no |
| Zhang et al.<sup>124</sup> | 2019 | name | mind | repeated | lab | night | 1 | yes | no | no |
| Klepel et al.<sup>128</sup> | 2019 | other | mind | single | lab | night | 1 | yes | no | no |
| Sikka et al.<sup>133</sup> | 2019 | alarm | dream | repeated | lab | night | >1 | yes | no | no |
| Sterpenich et al.<sup>102</sup> | 2020 | alarm | mind | repeated | lab | night | 1 | yes | yes | yes |
| Blanchette-Carrière et al.<sup>72</sup> | 2020 |  | dream | single | lab | nap | >1 | yes | no | no |
| Picard-Deland et al.<sup>109</sup> | 2020 | alarm | mind | single | lab | night | 1 | yes | no | no |
| Yu<sup>115</sup> | 2020 |  | mind | repeated | lab | night | >1 | yes | no | no |
| Vallat et al.<sup>117</sup> | 2020 | name | dream | single | lab | nap | 1 | yes | no | no |
| Martin et al.<sup>19</sup> | 2020 | name | mind | repeated | lab | night | >1 | yes | yes | yes |
| Scarpelli et al.<sup>134</sup> | 2020 | name | mind | single | lab | night | 1 | yes | no | no |
| Spanò et al.<sup>57</sup> | 2020 | alarm | mind | repeated | home | night | 1 | yes | no | no |
| Stephan et al.<sup>66</sup> | 2021 | alarm | mind | repeated | lab | night | >1 | yes | yes | yes |
| Picard-Deland et al.<sup>61</sup> | 2021 | alarm | dream | single | lab | nap | 1 | yes | no | no |
| Pires et al.<sup>129</sup> | 2021 |  | dream | repeated | lab | night | >1 | yes | no | no |
| Picard-Deland et al.<sup>131</sup> | 2021 | alarm | mind | repeated | lab |  |  | yes | yes | yes |
| Aamodt et al.<sup>42</sup> | 2021 | alarm | mind | repeated | lab | nap | 1 | yes | yes | yes |
| Wamsley<sup>132</sup> | 2022 | name | mind | repeated | lab | night | 1 | yes | no | no |
| Picard-Deland et al.<sup>58</sup> | 2023 | name | mind | repeated | lab | night | 1 | yes | no | no |
| Juan et al.<sup>75</sup> | 2023 | alarm | mind | repeated | lab | night | 1 | yes | yes | yes |
| Sebastiani et al.<sup>49</sup> | 2023 | other | dream | single | lab | nap | 1 | yes | no | no |
| Aamodt et al.<sup>43</sup> | 2023 | alarm | mind | repeated | lab | night | 1 | yes | yes | yes |

**Notes:** The awakening method 'name' means calling the participant by name, the awakening
method 'alarm' stands for waking up with an alarm/buzzer/sound/tone, question type 'mind'
corresponds to 'What went through your mind?', and the question type 'dream' to 'What did you
dream?'. In general, the label 'other' indicates a category that does not fall into the main
categories. Numbers in superscript in the publication column correspond to the references.

The question type ‘What went through your mind before the awakening?’ was asked in 75% of the cases, ‘What did you dream?’ in 19.1% and 5.9% were other questions. The awakenings were repeated within each recording in 80.9% of the studies, whereas single awakenings (17.6%) and a mixture of repeated and single awakenings (1.5%) were less frequent. The sleep environment was primarily the laboratory setting (91.2%), in comparison to at home (7.4%) and both (1.5%). Multiple and single day studies were 44.8% and 55.2% respectively. Most studies investigated night sleep (79.1%); others involved naps (20.9%).

The overall mean of study-based mean ages was 26.9 years with a standard deviation of 11.5 years. The overall mean percentage of females was 57% ± 21.2%. Furthermore, there were on average 27.2 ± 46.4 unique participants in a study. The overall mean of the total number of awakenings per study was 92.1 ± 135.8, amounting to roughly 3.4 ± 2.9 awakenings per person. The reviewed studies reveal an average percentage of experiences with recall of 59.5% in NREM sleep (considering only N1, N2 or N3 yields an average of 84.9%, 53.2% and 50.8% respectively), 83.4% in REM sleep and 95.3% in W. Experiences without recall occur on average 23.8% in NREM sleep (N1 = 10%, N2 = 22% and N3 = 28.9%), 7.1% in REM and 6.6% in W. Getting no report has values of 28% in NREM (N1 = 2%, N2 = 35.9% and N3 = 21.1%), 13.2% in REM and 3.6% in W.

### Experience models: Review

The experience with recall model revealed significant differences in NREM – REM (estimate = - 28%, p-value corrected < 0.001, Cohen’s d = 1.43) and NREM – W (-36.7%, p = 0.006, d = - 2.15), see Table 2 and Figure 1. For contextual variables the awakening method calling by name – alarm/buzzer/sound/tone (+10.1%, p = 0.046, d = 0.59), the study setting laboratory – at home (-14.4%, p = 0.046, d = -0.84) and the number of study days multiple – single (-8.9%, p = 0.021, d = -0.52) were significant, see Figure 2. The average number of awakenings per participant was not significant, see supplement Table S3. There were non-significant trends in the mean age and percentage females, but with a small effect size, see Table 2. Between N2 and N3 there were no significant differences. There was only one significant interaction between sleep stage and awakening method, where calling by name gives higher values in NREM than in REM, see Figure 2 and supplement Table S2.

**Figure 1.**
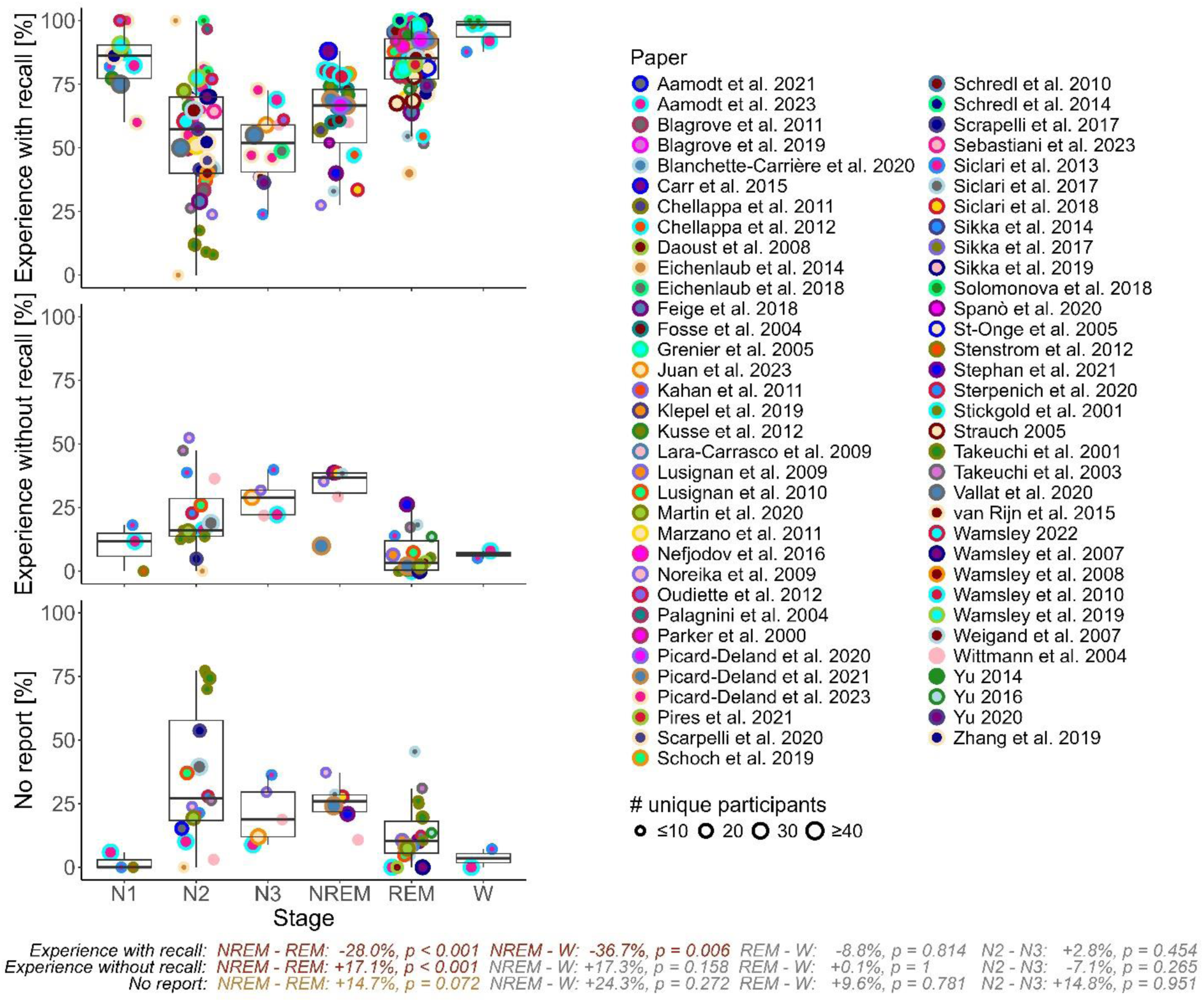
Percentages of experience type per sleep stage. All reviewed 69 studies and their percentages of experience with recall (top), experience without recall (middle) and no report (bottom) are shown. Some studies use the NREM, REM and W categorization, while others further looked into N1, N2 and/or N3. Text at the bottom indicates the linear model estimates on differences in sleep stages as well as their Benjamini-Hochberg corrected p-value. The size of the dots corresponds to the number of unique participants in a given study. There is a unique combination of dot fill and border color for each study. **Notes:** Here the individual studies’ categories NREM, N1, N2 and N3 were combined into one single NREM stage. Significant values (p ≤ 0.05) are in red, trends (0.05 < p ≤ 0.1) in orange. **Abbreviations:** Rapid eye movement (REM), non-REM (NREM), NREM stage 2 (N2), NREM stage 3 (N3), wake (W).

**Figure 2.**
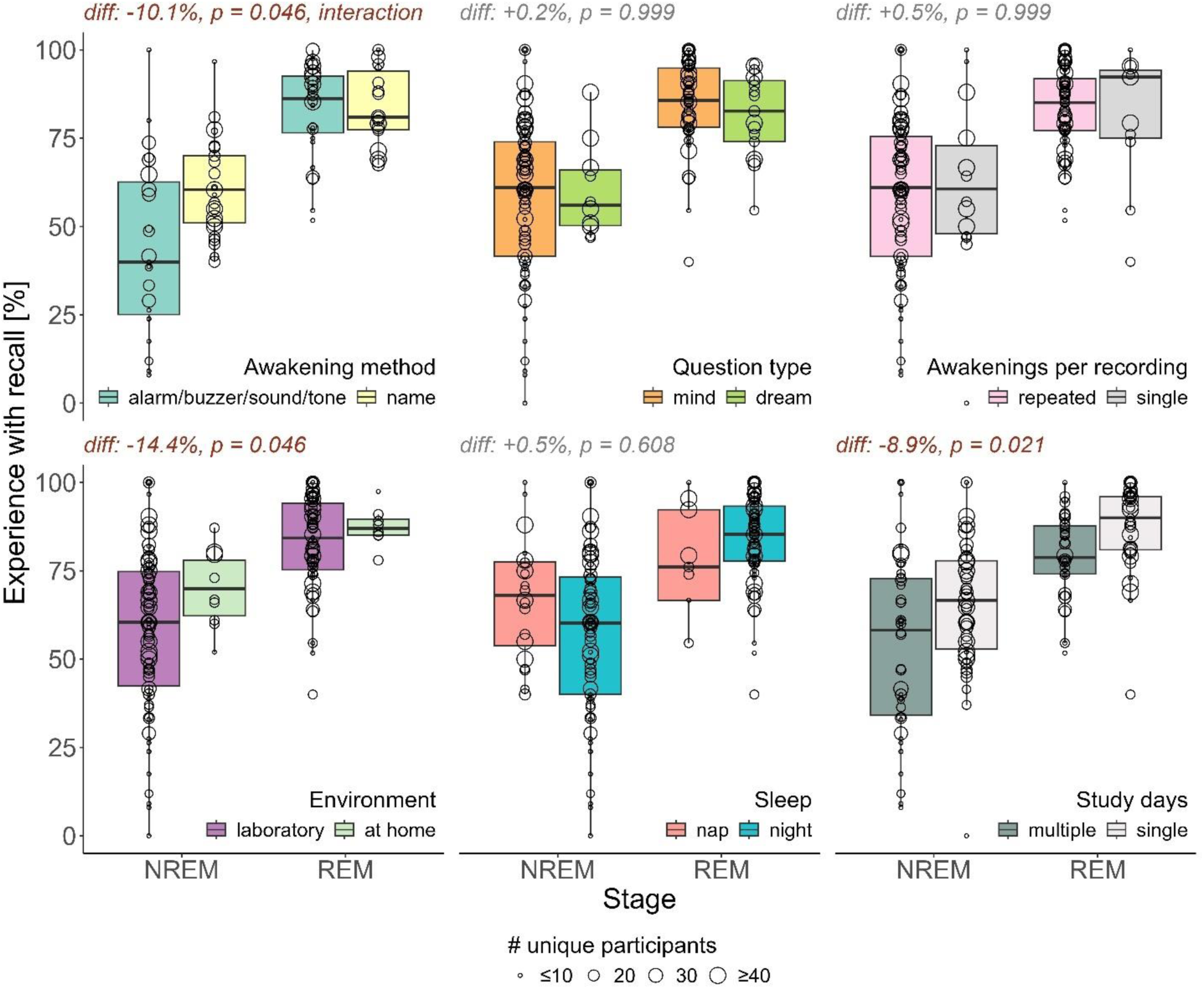
Boxplot of contextual variables. Here boxplots for the 69 reviewed studies with outcome percentage of experience with recall are shown. Furthermore, the estimated differences (diff) and Benjamini-Hochberg corrected p-values (p) from the linear model with intercept, sleep stage, awakening method, question type, mean age of participants, percentage of females, awakenings per recording, sleep environment, study days and sleep type are reported. The size of the dots corresponds to the number of unique participants in a given study. **Notes:** Here the individual studies’ categories NREM, N1, N2 and N3 were combined into one single NREM stage. The awakening method ‘name’ means calling the participant by name, question type ‘mind’ corresponds to ‘What went through your mind?’, and the question type ‘dream’ to ‘What did you dream?’. Significant values (p ≤ 0.05) are in red. **Abbreviations:** Rapid eye movement (REM), non-REM (NREM), difference (diff), number of (#).

**Table 2.**
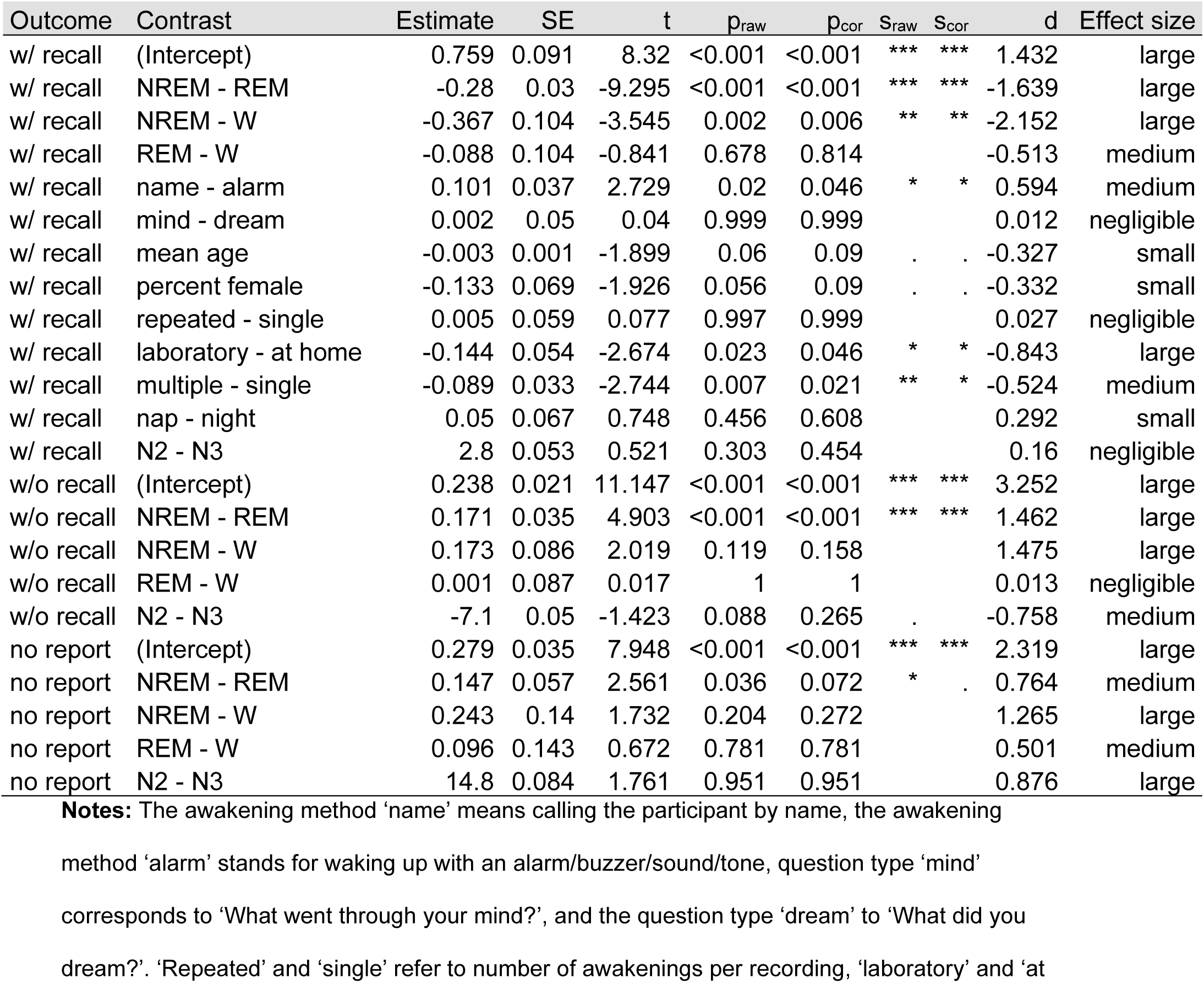
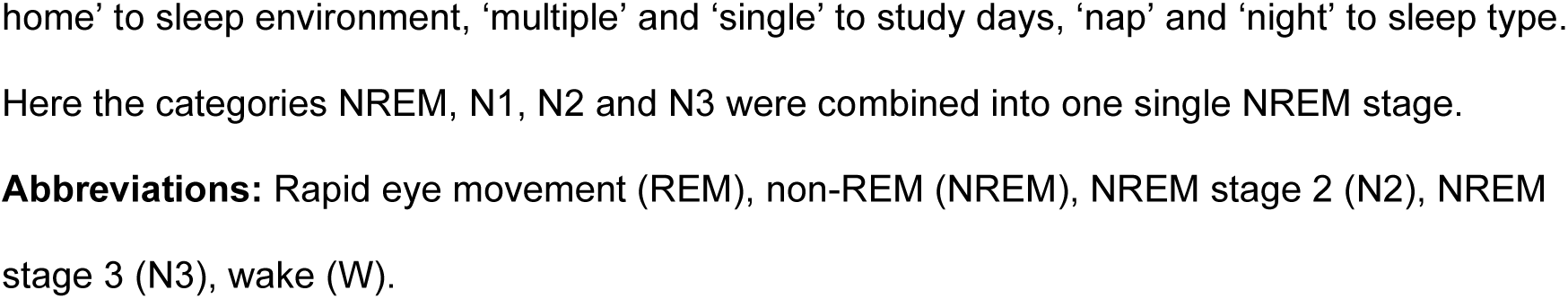
Models on awakening procedures of reviewed studies. Here linear models for the 69 reviewed studies with outcomes percentage of experience with recall (w/ recall), experience without recall (w/o recall) and no report are shown. The model with recall is the most extensive with intercept, sleep stage, awakening method, question type, mean age of participants, percentage of females, awakenings per recording, sleep environment, study days and sleep type. Whereas the model without recall and no report only include sleep stage. The N2 to N3 comparison employs a complementary t-test on sleep depth. For each test the model estimate and its according standard error (SE), t-value (t) and p-value uncorrected (p_raw_) as well as corresponding significance stars (s_raw,_ ‘***’ ↔ p < 0.001, ‘**’ ↔ 0.001 ≤ p < 0.01, ‘*’ ↔ 0.01 ≤ p < 0.05, ‘.’ ↔ 0.05 ≤ p < 0.1, ‘ ‘ ↔ p ≥ 0.1) and Cohen’s d values (d) with their effect size category (‘negligible’ ↔ |d| < 0.2, ‘small’ ↔ 0.2 ≤ |d| < 0.5, ‘medium’ ↔ 0.5 ≤ |d| < 0.8, ‘large’ ↔ |d| ≥ 0.8) are reported. Multiple testing corrections based on Benjamini-Hochberg are provided as corrected p-values (p_cor_) with their significance stars (s_cor_, for uncorrected values s_raw_).

For the model on experience without recall, there was a significant difference between NREM – REM (+17.1%, p < 0.001, d = 1.46). Other stage differences were not significant, even though NREM – W displays a similar Cohen’s d value of 1.48, see Table 2. No significant difference between N2 and N3 was found.

The no report model displayed only a trend between NREM – REM with a medium effect size, see Table 2. No significant difference between N2 and N3 was found.

Additional supplemental analyses on the reviewed studies revealed no significant effect of year of publication or number of participants for all experience types, but studies describing the awakening as gentle had significantly higher experience with recall percentages (+7%, p = 0.033), see supplement Figure S1 and S2.

### Participant-level: DREAM database

Overall, the DREAM database included 478 participants from 18 studies. For the analysis of the DREAM database with participant-based values, the model of experience with recall displayed significant differences for N2 – REM (p < 0.001, d = -0.18), N2 – W (p = 0.003, d = -0.18), N3 – REM (p < 0.001, d = -0.22) and N3 – W (p = 0.004, d = -0.22). Other stage comparisons were not significant, nor were there any trends. The full results can be found in Table 3 and Figure 3B.

**Figure 3.**
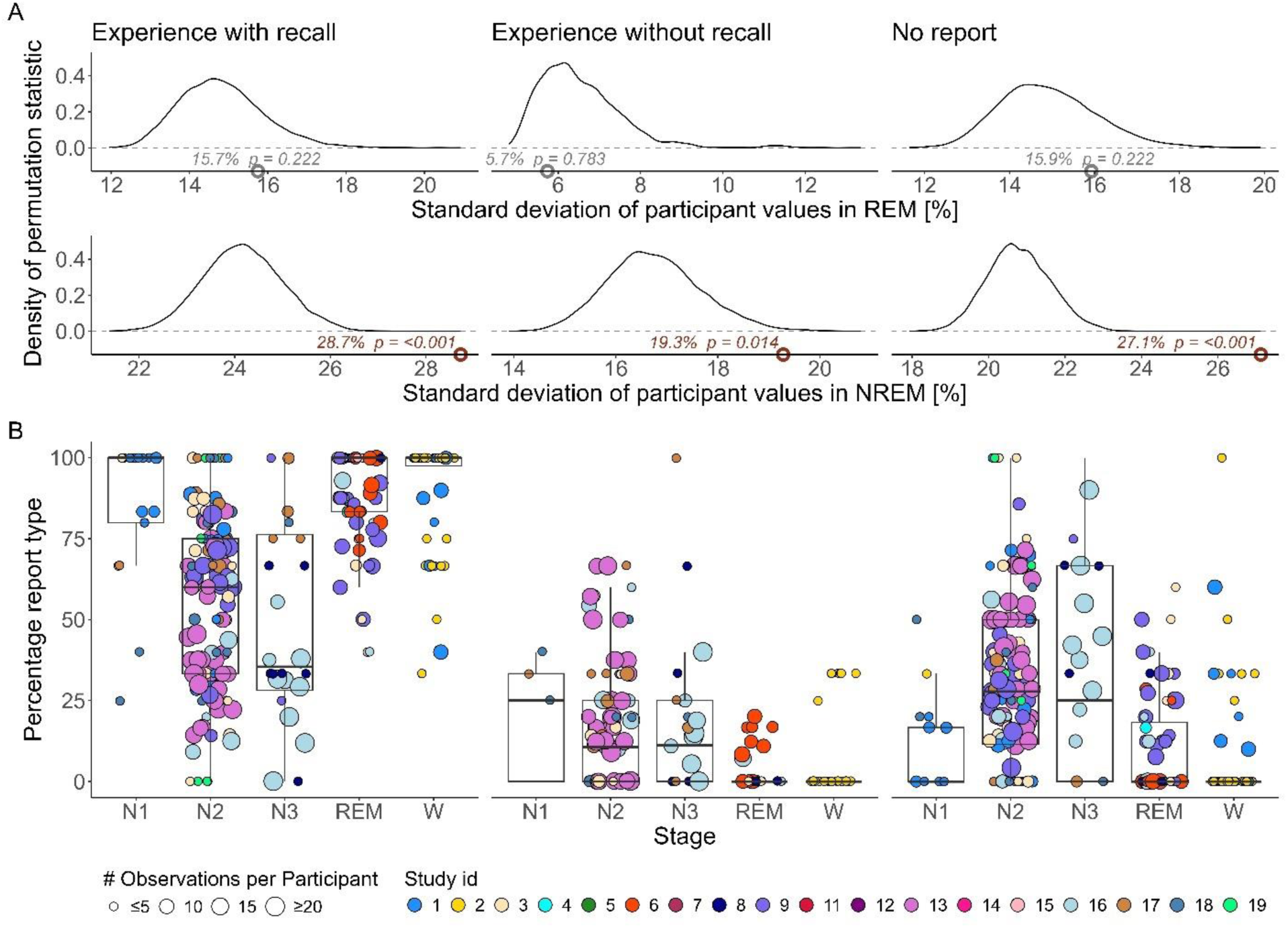
Effects of participant characteristics and percentages of experience per sleep stage in the DREAM database. Panel A shows the density of the permutation statistic per report type (columns experience with recall, experience without recall and no report) and sleep stage (rows REM and NREM) for the DREAM database. The permutation statistic captures influences from participant characteristics that go beyond age, gender and study design as measured by the standard deviation of participant percentage values. The observed values are indicated as dots with their according value and Benjamini-Hochberg corrected p-value (p) as text. Significantly large observed values thus indicate a higher-than-chance variability in participant percentages. In Panel B the participant-based percentages per report type (columns experience with recall, experience without recall and no report) and sleep stage (x-axis N1, N2, N3, REM and W) are displayed. **Notes:** In Panel A the individual studies’ categories NREM, N1, N2 and N3 were combined into one single NREM stage, permutation statistics only based on N2 can be found in the supplement. Significant values (p ≤ 0.05) are in red. The size of the dots corresponds to the number of unique awakenings per participant. The fill color of the dots indicate different studies available in the DREAM database. **Abbreviations:** Rapid eye movement (REM), non-REM (NREM), percent (%), number of (#).

**Table 3.**
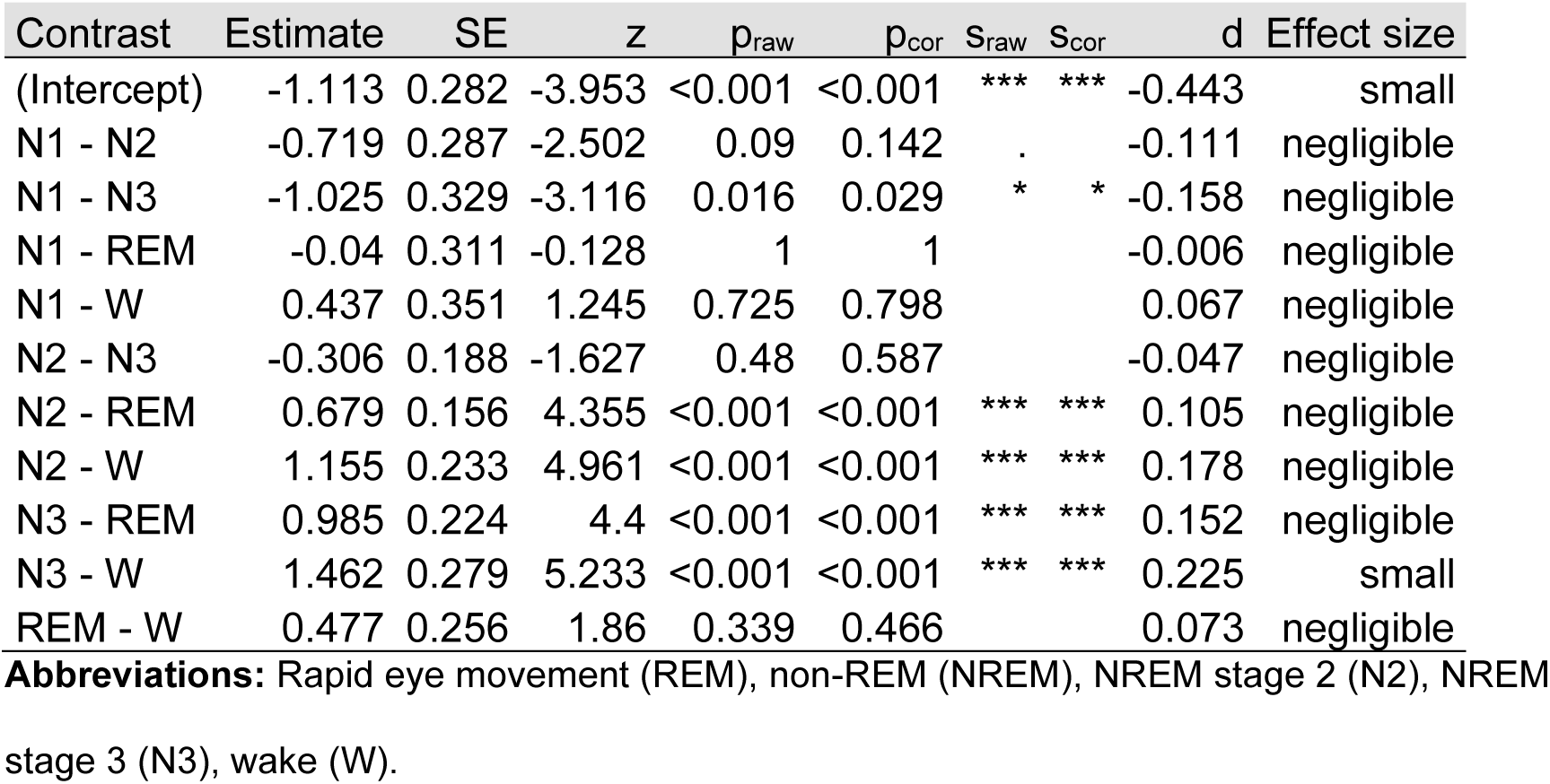
Models on sleep stage differences in the DREAM database. Here, the linear model for the DREAM database with the outcome variable percentage of experience with recall is presented. This percentage is calculated for each participant individually, provided they have three or more data points. For each test the model estimate and its according standard error (SE), z-value (z) and p-value uncorrected (p_raw_) as well as corresponding significance stars (s_raw,_ ‘***’ ↔ p < 0.001, ‘**’ ↔ 0.001 ≤ p < 0.01, ‘*’ ↔ 0.01 ≤ p < 0.05, ‘.’ ↔ 0.05 ≤ p < 0.1, ‘ ‘ ↔ p ≥ 0.1) and Cohen’s d values (d) with their effect size category (‘negligible’ ↔ |d| < 0.2, ‘small’ ↔ 0.2 ≤ |d| < 0.5, ‘medium’ ↔ 0.5 ≤ |d| < 0.8, ‘large’ ↔ |d| ≥ 0.8) are reported. Multiple testing corrections based on Benjamini-Hochberg are provided as corrected p-values (p_cor_) with their significance stars (s_cor_, with the same categorization as s_raw_).

For the DREAM database model on experience without recall, there was only a significant difference between N2 – REM (p = 0.017, d = 0.11). Furthermore, the model on no reports showed significant differences between N2 – REM (p < 0.001, d = -0.11), N2 – W (p < 0.001, d = -0.18), N3 – N1 (p = 0.029, d = -0.158), N3 – REM (p < 0.001, d = -0.15) and N3 – W (p < 0.001, d = -0.23). The results can be found in Table 3 and Figure 3B.

In NREM 151 unique participants from 9 studies with more or equal than 3 awakenings remained. For REM there were 70 unique participants from 7 studies with more or equal than 3 awakenings. The permutation tests to assess participant characteristics within the DREAM database were significant for NREM but not REM. Similarly strong significances can be found for N2, see supplement Figure S3. Figure 3A displays the test statistic distributions and observed values.

Experience with recall has an observed standard deviation of 28.7% (p < 0.001), experience without recall 19.3% (p = 0.014) and no report 27.1% (p < 0.001), see Figure 3A.

## Discussion

A review by Nielsen on awakening studies conducted before 2000 described an increase in NREM experiences since the 1950s.^12^ Our review with studies between 2000 and 2024 revealed that awakenings from NREM sleep lead on average to 60% experiences with recall, compared to 83% in REM sleep, aligning well with Nielsen’s findings of studies in the 1990s. This indicates that the observed increase has stabilized over the past 30 years, a conclusion further supported by non-significant yearly trends in our data (see Supplement Figure S1). Including the category of experiences without recall, which Nielsen’s review did not consider, revealed that these so called ‘white dreams’ predominantly occur during N2 and N3 sleep stages. Furthermore, getting no report occurs rarely, but on average twice as often in N2 and N3 than in REM. The approximate 30% decrease in experience with recall in NREM compared to either REM or W is evenly distributed among increases in experiences without recall and getting no report.

At first glance these results might suggest that N2 and N3, characterized by sleep spindles, K-complexes and slow waves, are somewhat more unconscious states than N1, REM and W. This link between slow waves and loss of consciousness has been put forth since the 1950s.^136^ One recent study found that boosting slow waves with closed loop auditory stimulation increased the experience without recall and no report conditions.

But upon closer examination of our study- and participant-level analyses we find no significant differences between N2 and N3 sleep for each experience condition. Getting no report is even estimated to be slightly lower in N3 than N2. If slow waves are causally implicated in suppressing the generation of experience this result seems odd and warrants further explanation. One such reason could be that most studies focused on N2 instead of N3 which leads to reductions in statistical power. Auditory thresholds for awakening participants have been found to be highest in N3.^137^ Hence, difficulty in awakening participants from N3 sleep could result in missing null-reports and skewed percentages. However, this explanation appears inconsistent, as REM sleep has comparably high arousal thresholds—lower than N3 but higher than N2^137^—while still permitting substantial dream recall. Another potential explanation is that unmeasured study effects outweigh the influence of sleep depth. For example studies capturing both N2 and N3 experiential reports often report slightly higher null report rates in N3 than N2.^43–45,65^ Nonetheless, the reported increases are small, indicating that sleep depth, quantified here as N2 and N3, is not a strong predictor for experiences during sleep. It was also suggested that slow waves and high-frequency components occurring over posterior regions are important.^20^ But an explanation is warranted why these types of oscillations need to occur, as suggested by our results, with a similar frequency in N2 and N3.^75^ On the other hand a review on studies correlating delta activity with dreaming was inconclusive, where only about half of the studies report an association.^40^ Furthermore, interventional studies showing decreases in experience with recall cannot conclusively prove that experience generation is suppressed,^138^ as interventions may hinder dream encoding or retrieval.^81,139^ Thus, the non-significant difference between N2 and N3 could also reflect an actuality and align with observations indicating that experiences during anesthesia are not related to the depth of anesthesia.^140^ It could rather be that increases in null reports in N2 and N3 compared to N1, REM, and W in part reflect a reduced saliency of NREM experiences.^141^ Experiences in N2 and N3 might be more subtle, including experiences lacking content or self-awareness,^142–145^ and are thus not prominent enough to be kept in memory.

Another unexpected finding was that the question type, even though slightly lower recall for asking about dreams compared to mental activity (Figure 2), was not significant. Some of the reviewed studies may have instructed participants to report dreams, explaining that this includes any mental activity, but did not mention this clarification in the paper, thereby masking any effects. It could also be that the questions ‘What went through your mind?’ and ‘What did you dream?’ yield similar responses. Both wordings may imply a limited focus on thinking and imagery and miss more subtle experiences, like feeling a sense of calm.^142,146,147^ As mentioned above, states of pure awareness during sleep, lacking rich content, have been observed.^145,148^ However, participants might not recognize these as states of consciousness, or if they do, fail to categorize them as a dream or mental activity. Thus, refining the questions being asked remains relevant for future research.

One of the main findings was that the awakening procedure is associated with dream recall rates. Generally, calling the participant by name corresponds to increases in experience with recall rates compared to using an alarm, especially when the awakenings were described as gentle. This finding is in line with the interference hypothesis,^149^ stating that dream recall can be more difficult in the presence of distractors such as an alarm clock.^150^ Greater reductions in cortisol during retrieval, likely to be observed during less stressful awakenings, was found to benefit episodic memory recall.^151^ Furthermore, it is known that the brain state from which one is awakened can persist for some time.^117,152,153^ If the awakening is gentle, the participant may remain longer in a state resembling the prior sleep experience, thereby improving recall due to the state-dependent nature of memory.^154,155^ Additionally, successful long-term memory encoding has been found to require a 2 minutes awakening^139^, emphasizing the need for some uninterrupted time for retrieval^34^ or memorization.^150^ Future studies should thus investigate the effect of awakening protocols more thoroughly.

Furthermore, the study’s environment may influence dream recall, as suggested by the results. Home studies yielded higher recall rates, which might be due to the participant being in a familiar sleep environment. In the laboratory, first night effects of dream recall could be present, most noticeable in studies with no adaptation nights or nonconsecutive experimental nights.^156^ Another explanation could be that home studies were often based on algorithms, which can lead to misclassifying sleep stages and in turn affect recall rates. Nonetheless, two of the studies conducted polysomnography at home.

Another effect related to the study design is that studies extending over several days display reduced dream recall. This effect might reflect poor adherence to the study protocol, likely caused by the tiring nature of undergoing multiple days of awakenings. No such effects were observed for multiple awakenings per recording. Hence, many awakenings in a single night might be less tiring than multiple days of awakenings. Since we did not quantify the exact number of awakenings for each participant per recording, excessive awakenings impairing dream recall cannot be ruled out. Lastly, another important finding from this study suggests the presence of participant characteristics that cannot be attributed to age, gender or differences in study design. These results might be taken as evidence for people having different levels of consciousness during sleep, but they could just as well point to dream encoding or retrieval.^157^ Variables affecting morning dream recall, such as openness to experience^82–86^ or the intention to remember a dream^150^ are likely candidates to also influence experience sampling from awakenings. Future studies are needed to pinpoint the sources of these effects.

This review is subject to certain limitations. First, likely not all awakening studies since 2000 have been identified, potentially influencing the results. Nonetheless, numerous studies from diverse research groups have been included, ensuring a more balanced outcome. Second, only a few contextual variables were readily available for most studies, hence limiting some questions to be addressed in this review. Third, the analyses are correlational and not suited for establishing definitive causal relationships. However, these analyses paint a coherent picture and can inform strategies for optimizing future awakening studies. Last, this review and analysis, as well as current awakening studies, do not provide conclusive evidence for the degree to which the null reports reflect the absence of phenomenal experience, or whether they are due to other effects.^157^ Our results only suggest, through the observed significant contextual and participant variables, that some of the null reports are not due to being conscious or not. This needs to be further investigated and if possible quantified.

## Conclusion

Our review of 69 awakening studies and the analysis of the DREAM database indicate that contextual variables and participant characteristics play an important role in experience recall rates. In particular, the method of awakening, the sleep environment, and participant characteristics beyond age, sex and study design are crucial factors. The awakening process and interindividual differences are especially important during NREM sleep, a state of fleeting memory with perhaps subtle experiences. To find meaningful neural correlates of consciousness, one requires good quality data on both the neural and subjective side. Hence, capturing the experiential aspect of reality is crucial and requires further investigation.

With rigorous subjective methodology in place, future research can employ machine learning to quantify the relationship between neuronal activity and experience, an area currently underexplored.^158^ Interpretable machine learning models provide insights into the aspects a model uses for decision making^159–161^ and thus serve as investigative tools to discover neural aspects of consciousness. Given the plethora of philosophical positions on consciousness,^162^ and the potential compatibility of current neuroscientific evidence with non-physicalist theories,^163^ our search for neural correlates should remain as philosophically neutral as possible. This means in particular to avoid the assumption that there is a direct correspondence between objective and subjective measures from the outset, and instead to explore the extent of their association. To this end, one could introduce benchmarks based on the generalizability of state-of-the-art machine learning models to unseen participants.

Consciousness during sleep can also be studied more directly, for example through communicating with people in lucid dreams, ie states of consciousness where one is aware that one is dreaming while dreaming.^164^ Real-time communication between a dreamer and the external world is possible even in NREM sleep stages^165^ and might be extendable to dreamless awareness.^142,144,145^ This holds the potential for overcoming some limitations of retrospective reports. But even during direct communication with the dreamer, participant characteristics and the questions asked likely play a role in the responses one gets. Some people may be more familiar with certain states of being than others, or give skewed reports containing preconceived opinions or interpretations, rather than observing what is experienced in an impartial way.

To overcome this, one of the most promising avenues is using first-person approaches, such as the micro-phenomenological interview where individuals undergo detailed interviews that aim at enhancing precision of reports.^146,148^ The Indian tradition has developed rigorous subjectivity for millennia,^166,167^ and hence we need to engage in a cross-cultural dialogue.

Another development that lead to large numbers of recalled dreams, even from N3, trained the participants in the questionnaire beforehand and clarified any uncertainties.^43^ A different study instructed participants to lie calmly in bed after the awakening with some time to let the memories come back.^34^

All of these avenues might sharpen the subjective data used in awakening studies and consciousness research in general.

In conclusion, our review of 69 studies between 2000 and 2024 and analysis of the DREAM database, in which participants were woken up to collect experience reports, shows that the context of the awakenings, such as the awakening process, the sleep environment and participant characteristics influence dream recall rates. This highlights the need for dream research to refine how reports are collected and pay closer attention to the subjective realm, where consciousness is to be found. We ourselves are the most direct sensors of consciousness, not any devices or algorithms. By refining our own “measurement instrument”, we may discover a wide range of states of consciousness in areas we previously thought none existed, enabling innovative ways of using these states and informing the treatment and care of disorders of consciousness.

## Acknowledgments

I wish to express my deepest gratitude to Prof. Dr. Hans-Peter Landolt for supporting me. He provided helpful feedback on the analyses and figures as well as pointing me towards some great papers. Furthermore, I would like to thank Prof. Dr. Peter Achermann and Laura Schnider for some insightful discussions.

Some sentences were revised for clarity using suggestions, alternative phrasings, or synonyms provided by the AI tools Microsoft Copilot (Microsoft Corporation, Redmond, Washington, USA, https://copilot.cloud.microsoft) and DeepL Write (DeepL GmbH, Cologne, Germany, https://www.deepl.com/en/write). Microsoft Copilot was additionally used, as indicated above, to find some studies with awakenings and experience collection. Furthermore, some R code modifications were assisted by Microsoft Copilot.

## Disclosure

The author reports no conflicts of interest. The work was supported by grant # 227100/Z/23/Z of the Wellcome Trust and Institutional funds of the University of Zurich.

The DREAM database is freely accessible (https://doi.org/10.26180/22133105) and the data set of the reviewed studies can be found as a supplemental excel sheet.

