## supplemental information for "We are the sensors of consciousness! A review and analysis on how awakenings during sleep influence dream recall"

Benjamin Stucky[1]

[1]Institute of Pharmacology and Toxicology, University of Zurich, Zurich, Zurich, Switzerland,

https://orcid.org/0000-0002-7425-2927

Correspondence: Benjamin Stucky

Institute of Pharmacology and Toxicology

University of Zurich

Winterthurerstrasse 190

CH-8057 Zurich

**Table S1** Included studies and their full characteristics

The table itself is too large to be displayed in the word document, but can be found as an Excel sheet (TableS1.xlsx). Here the 69 studies that were included in the review are listed. In addition to the information in Table 1, the mean age, female, number of unique participants and awakenings, as well as experience with recall, without recall and no report values are given. Here also the approximated values that were inferable from the paper or supplement of the paper are indicated.

**Notes:** The awakening method ‘name’ means calling the participant by name, the awakening method ‘alarm’ stands for waking up with an alarm/buzzer/sound/tone, question type ‘mind’ corresponds to ‘What went through your mind?’, and the question type ‘dream’ to ‘What did you dream?’. In general, the label ‘other’ indicates a category that does not fall into the main categories. Numbers in superscript in the publication column correspond to the actual references.

**Abbreviations:** Rapid eye movement (rem), non-REM (nrem), NREM stage 2 (N2), NREM stage 3 (N3).

**Table S2** Important interactions of awakening procedure variables in reviewed studies

A stepwise AIC model selection with both directions was carried out for a linear model where the main effects were always kept in and only the interaction terms with sleep stage were selected. This is an ANOVA table for the found linear model. Similar significance structure for main effects is observed in comparison to the main model from the paper. Only the interaction between sleep stage and awakening method was significantly selected, but a trend was present for the interaction of sleep stage and percent female.

| | Df | F value | P value |
|---|---|---|---|
| sleep stage | 2 | 48.7 | < 0.001 |
| awakening method | 2 | 12.4 | < 0.001 |
| question type | 2 | 0.1 | 0.895 |
| mean age | 1 | 2.3 | 0.133 |
| percent female | 1 | 7.2 | 0.008 |
| repeated | 2 | 0.6 | 0.567 |
| sleep environment | 2 | 3.0 | 0.053 |
| study days | 1 | 8.1 | 0.005 |

| | | | |
|---|---|---|---|
| sleep type | 1 | 0.6 | 0.431 |
| sleep stage:awakening method | 3 | 5.3 | 0.002 |
| sleep stage:percent female | 1 | 3.4 | 0.068 |

**Notes:** The symbol ‘:’ indicates interaction terms.

**Table S3** Average number of awakenings per participant

A linear model to predict percentage of experience with recall based on those 69 reviewed studies that had one study night was conducted with explanatory variables sleep stage, average number of awakenings per participant and their interaction. The according ANOVA table is provided. Except for sleep stage no significant result is observed for average number of awakenings per participant nor their interaction.

| | Df | F value | P value |
|---|---|---|---|
| sleep stage | 2 | 15.3 | <0.001 |
| average number of awakenings | 1 | 0.9 | 0.344 |
| sleep stage: average number of awakenings | 1 | 0.1 | 0.805 |

**Notes:** The symbol ‘:’ indicates interaction terms.

**Figure S1** Percentages of experience recall per year of publication.

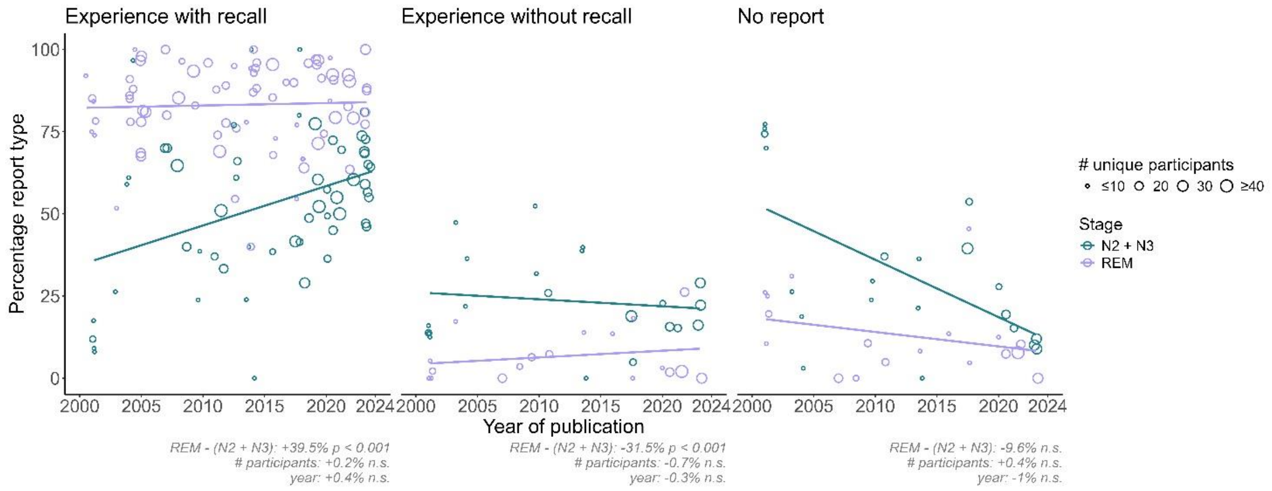


All the reviewed 69 studies and their according percentages of experience with recall, experience without recall and no report are shown per year of publication. Here, the focus lies on REM (purple) and combined N2 and N3 (green) sleep. The dots indicate number of unique participants in the study. Lines come from linear regressions with estimated percent changes per sleep stage, number of unique participants and year with their according uncorrected p-values are displayed. Expect for sleep stage differences not trend or interaction is significant. Visually, the trends for N2 and N3 approach the values for REM over time, but studies with more participants were also done more often towards the year 2024.

**Abbreviations:** Rapid eye movement (REM), non-REM (NREM), NREM stage 2 (N2), NREM stage 3 (N3), n.s. (not significant), # (number of).

**Figure S2** Gentle awakenings.

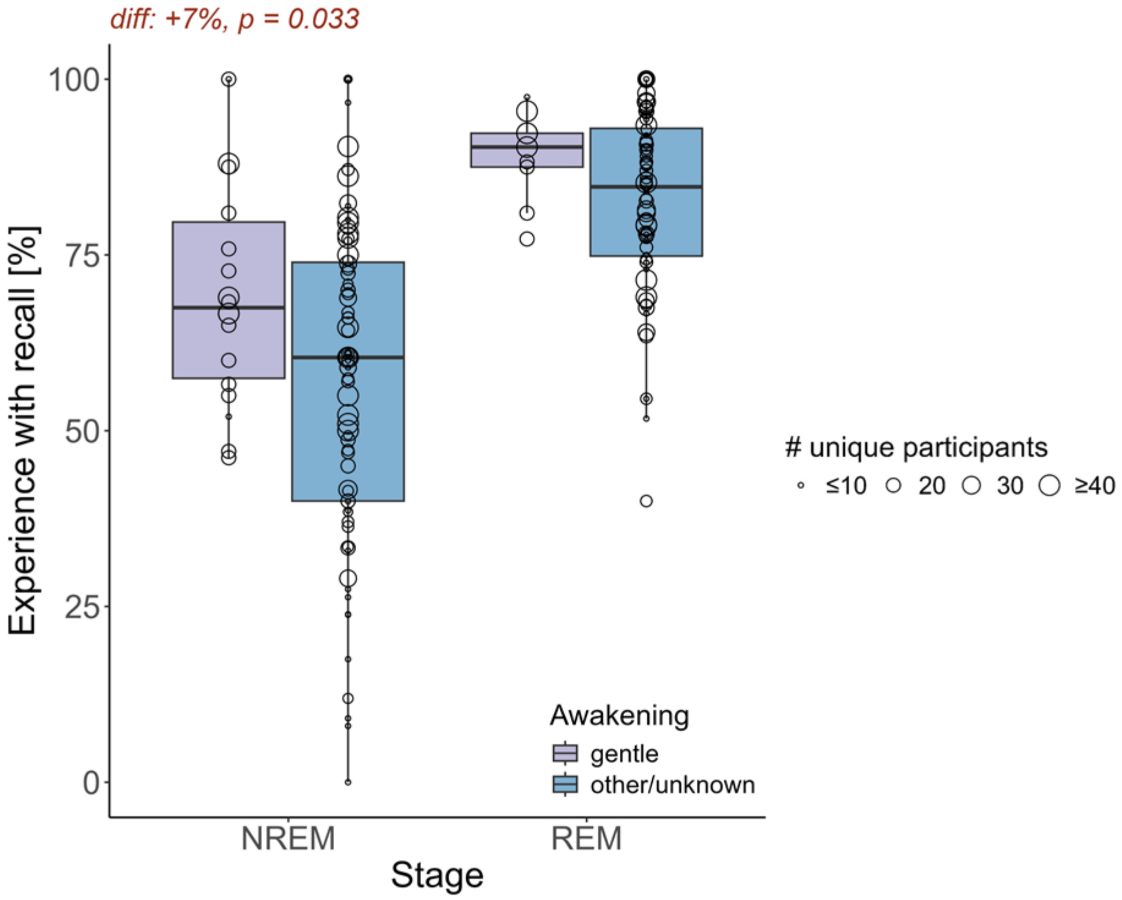


All the reviewed 69 studies were screened if they explicitly mention a gentle awakening (gentle) or not (other/unknown). This is a marginal t-test on the percentages of experience with recall. The estimated difference and uncorrected p-value are given. Gentle awakenings lead to 7% increases in recall rates, thus confirming the results were calling the participant's name, which possibly is a gentler awakening, displayed higher recall rates than awakening with an alarm. The size of the dots reflect the number of unique participants in each study.

**Abbreviations:** Rapid eye movement (REM), non-REM (NREM).

**Figure S3** Trait effects in DREAM database for N2.

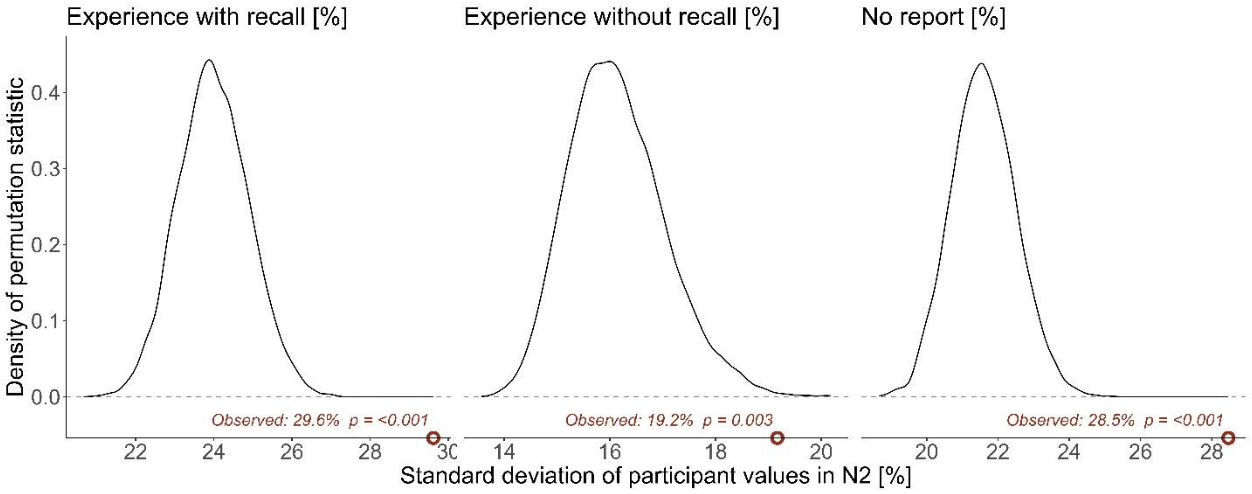


This figure shows the density of the permutation statistic per report type (columns experience with recall, experience without recall and no report) within the sleep stage N2. The permutation statistic captures traits that go beyond age, gender and study design as measured by the standard deviation of participant percentage values. The observed values are indicated as dots with their according value and uncorrected p-value (p) as text. Significantly large observed values thus indicate a higher-than-chance variability in participant percentages. The significance structure remains the same for N2 as compared to the combined NREM analysis in the paper.

**Notes:** Significant values ($p \leq 0.05$) are in red.

**Abbreviations:** Rapid eye movement (REM), non-REM (NREM), NREM stage 2 (N2), percent (%).
